# The loss of IL-31 signaling attenuates bleomycin-induced pulmonary fibrosis

**DOI:** 10.1101/2020.12.18.423450

**Authors:** Dan JK Yombo, Nishanth Gupta, Anil G. Jegga, Satish K Madala

## Abstract

Idiopathic Pulmonary Fibrosis (IPF) is a severe fibrotic lung disease characterized by excessive collagen deposition and progressive decline in lung function. Multiple Th2 T cell-derived cytokines including IL-4 and IL-13 have been shown to contribute to inflammation and fibrotic remodeling in multiple tissues. Interleukin-31 (IL-31) is a newly identified cytokine that is predominantly produced by CD4 TH2 T cells, but its signaling receptor called IL-31RA has been shown predominately expressed by non-hematopoietic cells. However, the potential role of the IL-31-IL31RA axis in pulmonary inflammation and fibrosis has remained largely unknown. To determine the role of IL-31 signaling in pulmonary fibrosis, wildtype, and IL-31RA knockout mice were treated with bleomycin and measured changes in collagen deposition and lung function. Notably, the loss of IL-31 signaling attenuated collagen deposition and lung function decline during bleomycin-induced pulmonary fibrosis. However, the loss of IL-31RA signaling did not affect inflammation in the lungs. The total lung transcriptome analysis showed a significant reduction in fibrosis-associated gene transcripts including ECM- and epithelial cell-associated gene networks. Furthermore, the lungs of IPF showed an elevated expression of IL-31 when compared to control subjects. In support, the percentage of IL-31 producing CD4^+^ T cells was greater in the lungs and PBMCs from IPF patients compared to healthy controls. Our findings suggest a pathogenic role for IL-31/IL-31RA signaling during bleomycin-induced pulmonary fibrosis. In summary, therapeutic targeting of the IL-31-IL-31RA axis could be beneficial in pulmonary fibrosis and has translational benefits specifically by preventing collagen deposition and improving lung function.

## Introduction

Pulmonary fibrosis is a chronic heterogeneous fibrotic lung disease characterized by an uncontrolled injury and repair process in the lung parenchyma. This process includes excessive deposition of collagen and extracellular matrix (ECM) protein components in the distal areas of the lung [1; 2]. Patients with severe fibrotic lung diseases including idiopathic pulmonary fibrosis (IPF) develop irreversible decline of lung function characterized by a decrease in forced expiratory volume in one second (FEV1) in part due to progressive accumulation of ECM producing myofibroblasts. Currently, FDA approved therapies to treat IPF, Nintedanib or Pirfenidone, delay the rate of lung function decline but do not reverse ongoing fibrosis or improve survival [3; 4]. While the etiology of IPF is not well established, pathways underlying the pathogenesis of pulmonary fibrosis have been identified [2; 5]. The development of fibrotic lesions, specifically, is regulated by signaling pathways that are driven by multiple pro-fibrotic growth factors, including TGF-β, TGFα, CTGF, and cytokines such as IL-1β, IL-17, IL-4, and IL-13 [6; 7; 8]. Th2 cytokines IL-4 and IL-13 have been found to play a major role in the pathophysiology of fibrotic diseases and are considered potential therapeutic targets in fibrotic diseases [8]. IL-13 signaling can induce activation of fibroblasts and macrophages to produce ECM and constitutive factors such as proline, which are required for the biogenesis of collagen. IL-4 and IL-13 signal through a type II IL-4 receptor alpha (IL-4Rα); this receptor is expressed in multiple cell types, including macrophages, epithelial cells, and fibroblasts [9].

Interleukin-31 (IL-31) is a recently described cytokine in the gp130/IL-6 cytokine family and is mainly produced by type 2 helper T cells. It plays a major role in the pathogenesis of chronic inflammatory diseases including atopic dermatitis (AD) or eczema [10; 11; 12]. IL-31 signaling is mediated through the heterodimeric receptor composed of the IL-31 receptor A (IL-31RA) and the Oncostatin M receptor β (OSMR β). IL-31RA is expressed by various cells including immune cells, epithelial cells, and fibroblasts; these cells can secrete pro-inflammatory cytokines and chemokines following stimulation by IL-31[11; 13; 14; 15; 16]. The role of IL-31 in the development of inflammatory diseases including AD has been described and potential therapeutic agents have been generated to target IL-31/IL-31RA signaling [10; 11; 17; 18]. Recent studies have shown elevated levels of IL-31 in patients with AD or asthma and levels have correlated with disease severity [19; 20]. In fibrotic diseases, several IL-6 family cytokines including IL6 and OSM have been shown to induce pulmonary fibrosis, but the contribution of IL-31/IL-31RA signaling in lung fibrosis has remained unexplored [20; 21]. Our previous work has depicted the effector mechanism of IL-4/IL-13 signaling in the pathogenesis of pulmonary fibrosis and the upregulation of IL-31RA via the IL-4Rα/STAT6 signaling axis [8; 22]. However, the role of IL-31/IL-31RA signaling in the pathogenesis of pulmonary fibrosis has not been definitively determined.

The current study aimed to determine the role of IL-31/IL-31RA signaling in the pathogenesis of pulmonary fibrosis using a bleomycin-induced lung fibrosis model and IL-31RA deficient mice. We observed a significant decrease in collagen deposition and the expression of profibrotic genes in the lungs of IL-31RA deficient mice compared to wildtype mice treated with bleomycin. Notably, the loss of IL-31RA signaling was sufficient to attenuate worsening lung function. Further, we observed increased expression of IL-31 in lung tissue and CD4-positive T cells of IPF compared to healthy subjects.

## Material and methods

### Mice

IL31RA knockout (IL-31RA^-/-^) mice and their littermates wildtype (WT) mice of C57BL/6 background were used in this study to investigate the role of IL-31RA signaling in the development of pulmonary fibrosis [11]. Both male and female mice at 10–18 weeks of age were used for all of the experiments and were housed in the Cincinnati Children’s Hospital Medical Center animal facility under conditions approved by the American Association for the Accreditation of Laboratory Animal Care. All mice were maintained under aseptic conditions and received sterile food and water. The experiments were approved by the Institutional Animal Care and Use Committee.

### Bleomycin-induced pulmonary fibrosis model

IL-31RA^-/-^ and littermate wildtype mice were intradermally injected with bleomycin to induce lung fibrosis as previously described [8; 23]. Briefly, mice were injected with saline or bleomycin (6 U/kg of body weight) in 50 μl saline, once a day and during 5 consecutive days per week, for a total of 4 weeks, to induce pulmonary fibrosis. Mice were euthanized on day 28 and lung samples were collected to assess fibrosis using biochemical and molecular methods.

### Histology and lung hydroxyproline

The lungs were fixed with 10% neutral formalin and paraffin-embedded. The five-micron thick lung sections were prepared and stained with Masson’s Trichome to evaluate the deposition of collagen in lung tissues, a key feature of the fibrotic process. The deposition of collagen was assessed by measuring hydroxyproline levels in lung lysates using a colorimetric detection assay as previously described [24]. In brief, the right lung lobes were collected, weighed, and hydrolyzed in 4 ml of 6 N HCl overnight at 110°C. Hydrolyzed samples were neutralized with 1 N NaOH. For colorization, chloramine-T (0.05 M) and the aldehyde-perchloric acid reagent were added and samples were placed in a hot water bath at 60°C for 25 min. The concentration of hydroxyproline in the samples was determined using the standard curve. The hydroxyproline level was normalized to the lung weight and expressed in comparison to the level measured in the saline-treated wildtype control group [8].

### Lung function measurements

Murine lung function was measured using a computerized FlexiVent system (Flexiware version 7.5, SCIREQ, Montreal Canada) as previously described [25; 26; 27]. Briefly, mice were anesthetized with Ketamine/Xylazine and an incision was performed on the anterior area of the neck to expose the trachea. The trachea was then cannulated with a 20G cannula and the mouse was connected to the FlexiVent system. Following deep inflation, three consecutive measures were collected for each mouse and the average value was used to measure resistance, elastance, and compliance.

### Immunohistochemistry

Formalin-fixed and paraffin-embedded human lung tissue sections of IPF (n=8) and healthy (n=7) controls were immunostained with antibodies against IL-31 as previously described [28; 29]. Briefly, rabbit anti-human IL-31 monoclonal antibody (R&D Systems) was used as the primary antibody (1:25 dilution). Goat anti-rabbit Ig was conjugated with a peroxidase enzyme to form a brown precipitate in the presence of hydrogen peroxide and DAB. Nuclei were stained in blue using Hematoxylin counterstain as described in previous reports [23; 30]. The staining of the lung section using control IgG showed no detectable immunostaining (data not shown). All images were obtained using a Leica DM2700 M bright-field microscope. High-magnification images (X40) were captured with a 3CCD color video camera. The number of IL-31-positive cells and total cells was counted using MetaMorph imaging software and expressed as the percentage of IL-31-positive cells in total cells in the field.

### RNA preparation and real-time PCR

Lungs were homogenized using Qiagen tissue homogenizer and beads (Qiagen), and total RNA was extracted from lung tissues using RNeasy mini kit (Qiagen Science, Valencia, CA) as previously described [10; 22]. Lung RNA samples were reverse transcribed into cDNA using Superscript III (Invitrogen), and the relative transcript expression of select genes was measured using SYBR Green PCR Master Mix (Applied Biosystems, Foster City, CA) and the CFX384 Touch Real-Time PCR detection system (Bio-Rad, Hercules, CA) as previously described [31]. Gene transcripts were normalized to the housekeeping gene hypoxanthine-guanine phosphoribosyltransferase (HPRT) or β actin and were expressed as the relative fold-induction change compared to the gene expression level of the control wildtype mice treated with saline solution. Data were analyzed with Bio-Rad CFX maestro software version 4.2(Bio-Rad Laboratories, Hercules, CA). The list of primers used for real-time PCR to analyze the relative expression of the genes is included in Supplementary Table 1.

### Whole lung transcriptome shotgun sequencing (RNAseq)

Total lung RNA samples were obtained from IL-31RA^-/-^ and wildtype mice treated with bleomycin as described above. Four samples were prepared from each experimental group and subjected to RNA sequencing using an Illumina HiSeq-1000 Sequencer (Illumina, San Diego, CA), as described previously [32]. A comparative analysis between groups was realized to identify genes with differential changes in expression following treatment with bleomycin or saline. Genes with statistically significant changes were selected based on a p-value cut-off of 0.05 and differential expression (defined as a 1.5-fold increase or decrease in expression). Heatmaps of genes with differential expression change were generated to highlight clusters of genes up-or downregulated in IL-31RA^-/-^ and wildtype mice treated with bleomycin. Functional enrichment analyses were performed using the ToppFun application of the ToppGene Suite [33]. The IL-31RA-dependent genes were also compared with differentially expressed genes (DEGs) from IPF patients (GSE53845) to identify common genes between IL-31RA regulated genes and IPF DEGs as described previously [34; 35].

### In vitro epithelial cell treatments

Human bronchial epithelial BEAS-2B cells were obtained from the American type culture collection (Manassas, VA) and cultured in DMEM medium supplemented with 10% fetal bovine serum (FBS) at 37°C and 5% CO2. To assess the effects of IL-31, airway epithelial cells were seeded in 12 or 24 well plates in low serum media (1%FBS) for 16 hrs. Under low serum conditions, cells were stimulated with media supplemented with increasing doses (0, 100, 250, or 500 ng/ml) of recombinant IL-31(R&D Systems) for 18 hours. Total RNA was extracted from the cultured cells using the Qiagen RNA shredder and the RNeasy mini kit (Qiagen). cDNA was prepared and applied for real-time RT-PCR to measure gene transcript levels as described above. Gene expression was normalized to the housekeeping gene human β-actin.

### Flow Cytometry

Peripheral blood mononuclear cells (PBMC) were isolated from peripheral venous blood of IPF patients (n=11) and healthy control donors (n=10) using Ficoll-paque density gradient-based separation with centrifugation (GE Healthcare, Uppsala, Sweden). Red blood cells were lysed using an ACK lysis buffer and cell viability was confirmed with trypan blue. Isolated PBMCs were frozen with DMSO and stored in liquid nitrogen until used. To measure the production of IL-31 and IL-4, PBMCs were thawed from liquid nitrogen and plated in a 96-well round-bottom plate (1 × 10^6^ cells/well) in RPMI 1640 medium supplemented with L-glutamine, antibiotics (penicillin/streptomycin), and 10% heat-inactivated fetal bovine serum (FBS). Cells then were allowed to rest overnight. PBMCs were stimulated with phorbol 12-myristate 13-acetate (PMA) 10 ng/ml with Ionomycin (1ug/ml), and Golgi stop (1:1500) for 5 hours to induce cytokine production. Control samples were cultured with PMA and ionomycin for the same period. Cells were stained with Zombie UV(BioLegend) for 30 min at room temperature in the dark to allow for discrimination between living and dead cells. Non-specific binding of fluorochromes was prevented by incubating cells with Human BD Fc Block™ (BD Biosciences), followed by surface marker staining with anti-CD3-V500 (UCHT1) and anti-CD4-APC(RPA-T4) (Bio Legend). Cells were then fixed and permeabilized using Cytofix/Cytoperm (BD Biosciences) and stained with antibodies against IL-31-PE (U26-947) (BD Biosciences) and IL-4 -PE-Cy7 (MP4-25D2) (BioLegend). Data were acquired using LSR Fortessa flow cytometer and analyzed using FlowJo 10 (FlowJo, Tree Star).

### Statistical analysis

Statistical analysis was performed using GraphPad PRISM 8 using a parametric student t-test to compare the mean between two groups. The mean between groups was compared using a one-way ANOVA test followed by Tukey’s post hoc analysis. Quantitative data were presented as mean ± SEM, and *p* < 0.05 was considered statistically significant.

## Results

### The loss of IL-31RA signaling attenuates collagen deposition and lung function decline

To determine whether the loss of IL-31RA has any effect on pulmonary fibrosis, IL-31RA knockout mice and their littermate wildtype mice were treated intradermally with bleomycin (to induce pulmonary fibrosis) or treated with saline as a non-fibrosis control. To assess collagen deposition, the lung sections were stained with Masson Trichrome. There were no significant differences in collagen staining between wildtype and IL-31RA knockout mice treated with saline. A significant increase in collagen deposition was observed in the lungs of both wildtype mice and IL-31RA knockout mice treated with bleomycin (Figure 1A). However, there was a modest decrease in collagen staining in mice deficient for IL-31RA compared to wildtype mice treated with bleomycin (Figure 1A). To further evaluate collagen deposition, the level of hydroxyproline was quantified in the lungs of IL-31RA knockout mice and wildtype mice treated with bleomycin or saline. Both wildtype and IL-31RA knockout mice treated with bleomycin had increased expression of lung hydroxyproline compared with saline-treated mice. However, there was a modest but significant reduction in lung hydroxyproline levels in IL-31RA knockout mice compared to wildtype mice treated with bleomycin (Figure 1B). Our published studies have demonstrated that repetitive bleomycin-induced fibrotic remodeling decreases lung function including resistance, elastance, and compliance [36; 37]. Therefore, we assessed whether the loss of IL-31RA signaling has any effect on the decline in lung function following repetitive bleomycin treatment. Consistent with previous findings, there was a significant increase in resistance, elastance, and compliance in wildtype mice treated with bleomycin compared to saline-treated wildtype or IL-31RA knockout mice (Figure 1C-1E). However, the decline in lung function was attenuated with the loss of IL-31RA signaling compared to wildtype mice treated with bleomycin (Figure 1C-1E). These findings suggest that the loss of IL-31RA has a partial protective effect against collagen deposition, and the subsequent decline in lung function, during bleomycin-induced pulmonary fibrosis.

**Figure 1.**
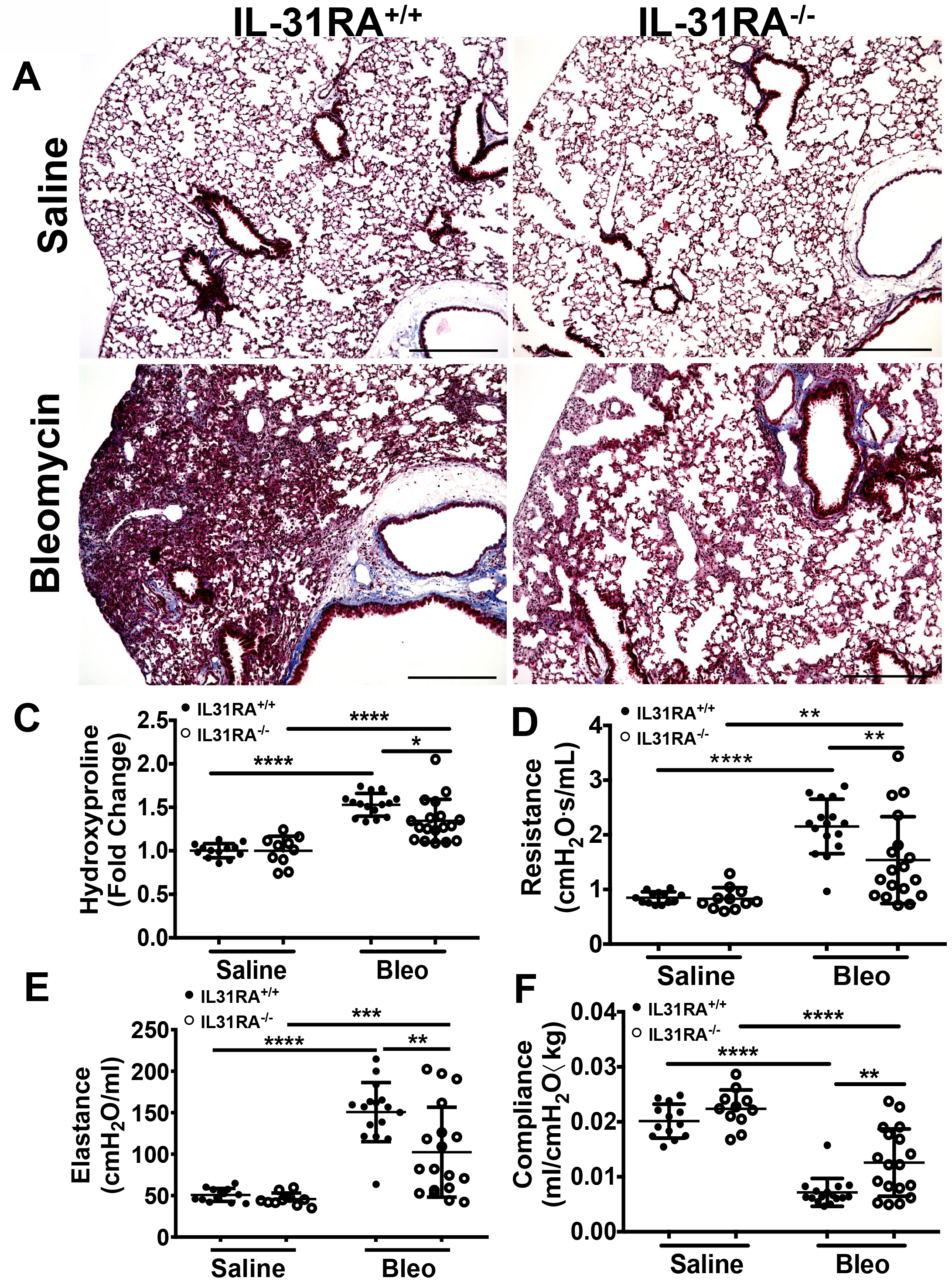
The loss of IL-31RA signaling attenuates collagen deposition and lung function decline during bleomycin-induced pulmonary fibrosis. wildtype (IL-31RA^+/+^) and IL-31RA knockout (IL31RA^-/-^) mice were treated intradermally with bleomycin or saline for four weeks to induce pulmonary fibrosis. (A) Lung sections of saline or bleomycin-treated mice were stained with Masson Trichrome. Scale bar 300 μm. (B) Total lung hydroxyproline was compared between IL-31RA^-/-^ and IL31RA^+/+^ mice treated with bleomycin or saline. (D-F) Lung function measurements including resistance, elastance and compliance were measured in all four groups using FlexiVent. Data are presented as means ± SEM. The above data is cumulative of two independent experiments (n=12-19/group). Statistical analysis was performed using one-way ANOVA with Tukey’s multiple comparisons test. **p < 0.01; ****p<0.001.

### The role of IL-31RA signaling in fibrosis-associated gene expression

To identify IL-31RA dependent gene expression and associated signaling pathways involved in bleomycin-induced pulmonary fibrosis, whole lung transcriptome analysis was performed using next-generation RNA-seq analysis in wildtype and IL-31RA knockout mice treated with bleomycin. The comparison of differentially expressed genes, with 1.5-fold up-or down-regulation in wildtype and IL31RA knockout lungs, highlighted two clusters of genes as illustrated in the heatmap (Figure 2). A total of 458 genes were differentially expressed in the lungs of IL-31RA knockout^-^ mice compared to wildtype mice treated with bleomycin. About 228 genes were up-regulated in IL-31RA knockout mice, and 230 genes were down-regulated, with the loss of IL-31RA signaling (Supplementary Table 2). Gene ontology enrichment analysis of down-regulated genes suggested IL-31RA is involved in the expression of several extracellular matrix (ECM)-associated genes (*Col1a1, Col3a, Fn1, Arg1, Timp1, Mmp13, Ereg, and IL-6)*, Protein G coupled receptor Gα(i) signaling (Ccl28, Rgs5, Rgs4, and Rgs16), Epithelium-associated genes (*Krt4, Krt5, Krt14, Krt20)*, Adenylate cyclase activity (*Calcb, Calcr*) and chemotaxis (*Ccl28, Mcp1, IL6*). Enrichment analysis of genes that were upregulated with the loss of IL-31RA suggested an increase in the expression of genes associated with the G protein-coupled receptor, and synaptic and calcium signaling.

**Figure 2.**
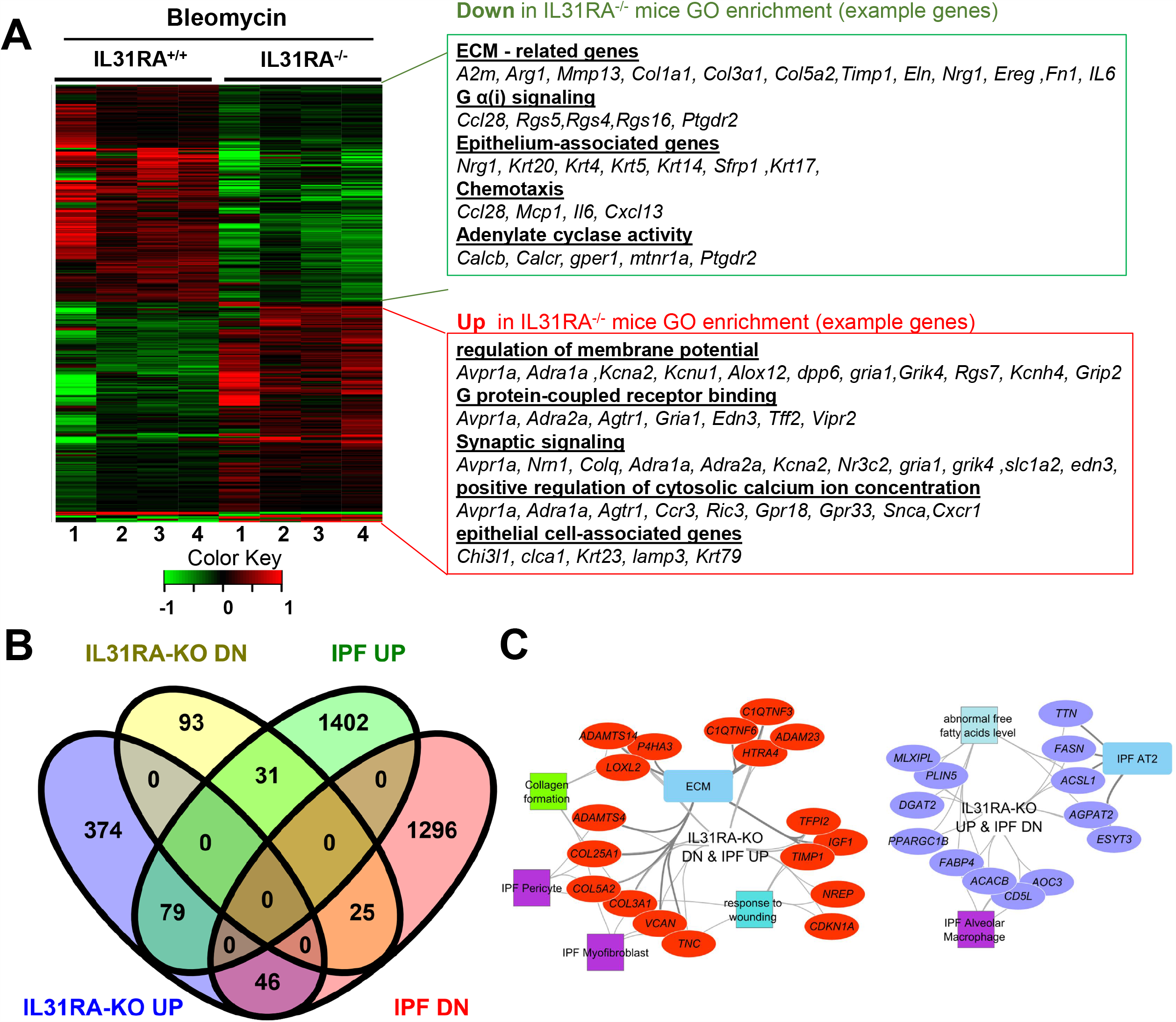
The IL-31-IL-31RA axis regulated gene networks during bleomycin-induced pulmonary fibrosis and IPF. Total lung RNA was isolated from bleomycin or saline treated mice and performed RNA-seq analysis. (A) Heat map shows two clusters of differentially expressed genes that were up- or down-regulated by at least 1.5-fold upon bleomycin treatment of genetic knockdown of IL-31RA compared to wildtype mice. Gene Ontology Enrichment analysis for IL-31RA-dependent gene transcripts was performed using ToppFun. (B) Venn diagram of IL31RA-regulated genes compared with IPF-differentially expressed genes in the lungs of IPF patients. (C) IL-31RA-driven gene networks that are activated in IPF were analyzed using ToppFun and visualized using Cytoscape. Red- and violet-colored oval shapes represent genes that are IL31RA-regulated and up-or down-regulated respectively in IPF lungs. Both square and rectangle shapes represent different enriched biological processes and cell types for the inversely correlated genes between the IL-31-IL-31RA axis and IPF.

Next, to identify the IL-31RA-dependent genes that are also dysregulated and relevant to IPF pathogenesis, differentially expressed genes in IPF lungs were obtained and compared to IL-31RA-dependent genes (Figure 2B). In total, 181 IPF differentially expressed genes were identified that were also dysregulated with the loss of IL-31RA (Figure 2B). This IL-31RA dependent gene list includes 31 genes that were up-regulated, and 46 down-regulated genes, in IPF lungs (Figure 2B and Supplementary Figure 1). We performed an enrichment analysis of these negatively correlated genes using the ToppFun application and visualized the gene networks using Cytoscape. As shown in Figure 2C, our gene functional enrichment analysis further supports alterations in fibroblast- and epithelial cell-associated genes and also a decrease in multiple collagen gene transcripts.

### The loss of IL-31RA signaling attenuates the expression of pro-fibrotic genes

To validate the findings using RNA-seq, fibrosis-associated gene transcripts were quantified in the total lung RNA of IL-31RA knockout and wildtype mice treated with bleomycin or saline. The major ECM gene transcripts including *Col1α, Col3α, and Fn1* were upregulated in lungs of wildtype mice treated with bleomycin compared to wildtype mice treated with saline. This increase in ECM gene expression was significantly attenuated in IL-31RA knockout mice compared to wildtype mice treated with bleomycin (Figure 3A). Similarly, genes associated with ECM production and remodeling were significantly decreased with the loss of IL-31RA; these genes included MMP13 and TIMP1, as well as IL-6 which increased in wildtype mice during bleomycin-induced pulmonary fibrosis (Figure 3B). The total lung RNA-seq analysis also suggests that altered expression of several epithelial cell-associated genes with the loss of IL-31RA during bleomycin-induced pulmonary fibrosis (Figure 2). To validate the differential expression of genes associated with epithelial cells, we measured transcript levels of several epithelium-associated genes including *Krt5*, and *Krt14*. As shown in Figure 4A, we observed a significant increase in the transcripts of Krt5, and Krt14 in wildtype mice treated with bleomycin compared to saline-treated wildtype mice. This increase in epithelial gene transcripts was attenuated with the loss of IL-31RA compared to wildtype mice treated with bleomycin (Figure 4A). These findings suggest a critical role for airway epithelium in the pathophysiology of IL-31RA-driven pulmonary fibrosis and decline in lung function. Therefore, we studied the effect of IL-31 on airway epithelial cells in the production of pro-fibrotic cytokines associated with pulmonary fibrosis. Bronchial epithelial cell line BEAS-2B, or primary bronchial epithelial cells, were stimulated with IL-31 cytokine and the transcript levels of *Mcp1* and *IL-6* were determined using RTPCR. The production of pro-fibrotic cytokines, including *Mcp1* and *IL-6*, increased with IL_31 stimulation in both BEAS-EB and HBEC cells (Fig. 4B & 4C). These new findings suggest that IL-31-driven signaling increases the production of pro-fibrotic cytokines by epithelial cells in the pathogenesis of pulmonary fibrosis.

**Figure 3.**
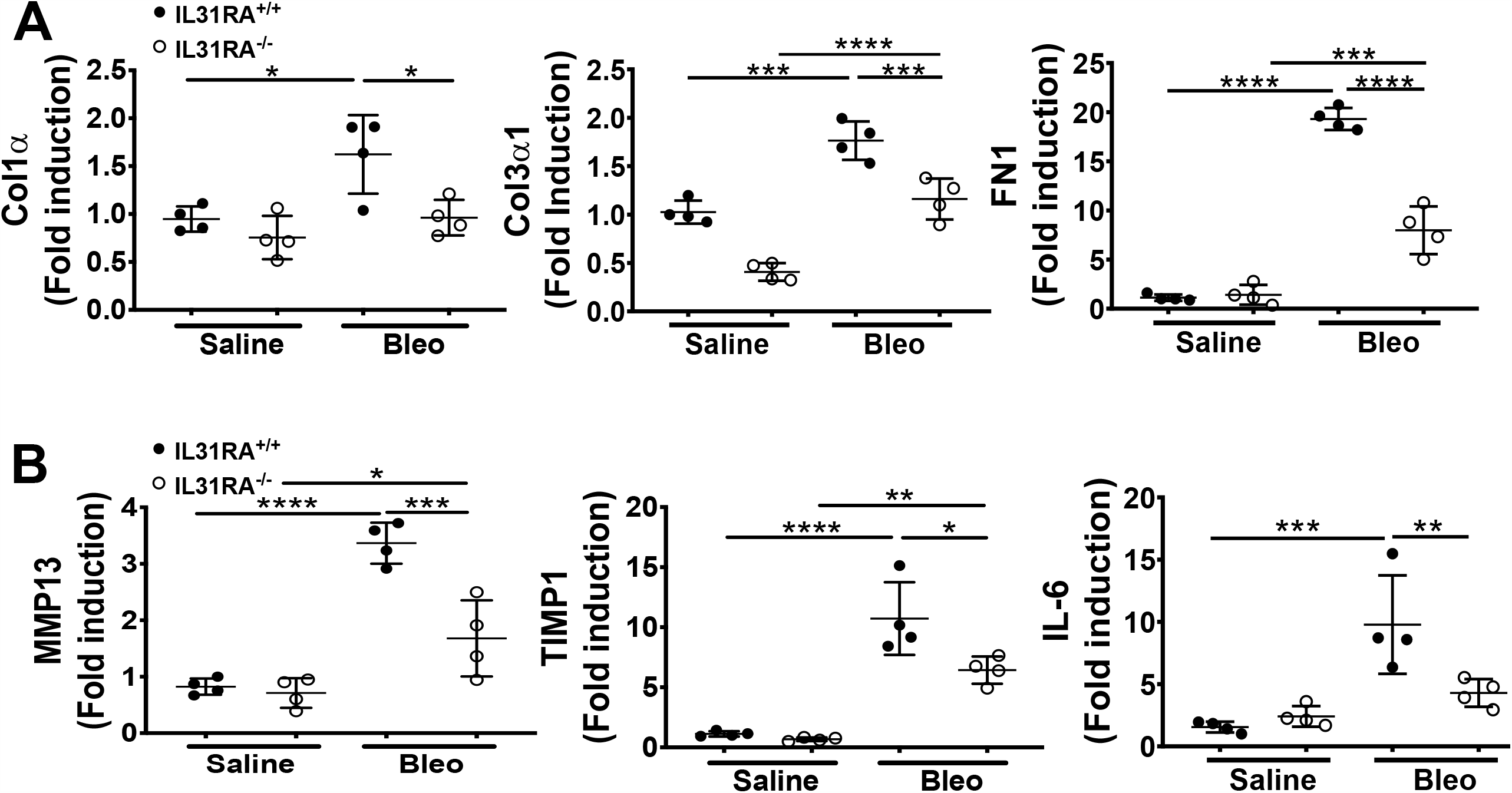
The loss of IL-31RA signaling attenuates the expression of fibrosis-associated genes during bleomycin-induced pulmonary fibrosis. Total lung RNA was isolated from wildtype and IL-31RA^-/-^ mice treated with saline and bleomycin and fibrosis-associated genes quantified using real-time RTPCR. (A) Quantification of ECM gene transcripts including Col1α, Col3α1, and Fn1. (B) Quantification of gene transcripts involved in ECM remodeling and production including Mmp13, Timp1 and Il6. Data presented as Mean ± SEM (n=4/group). Statistical analysis was performed using one-way ANOVA with Tukey’s multiple comparisons test * p<0.05; ** p<0.01; *** p< 0.005; **** p<0.001.

**Figure 4:**
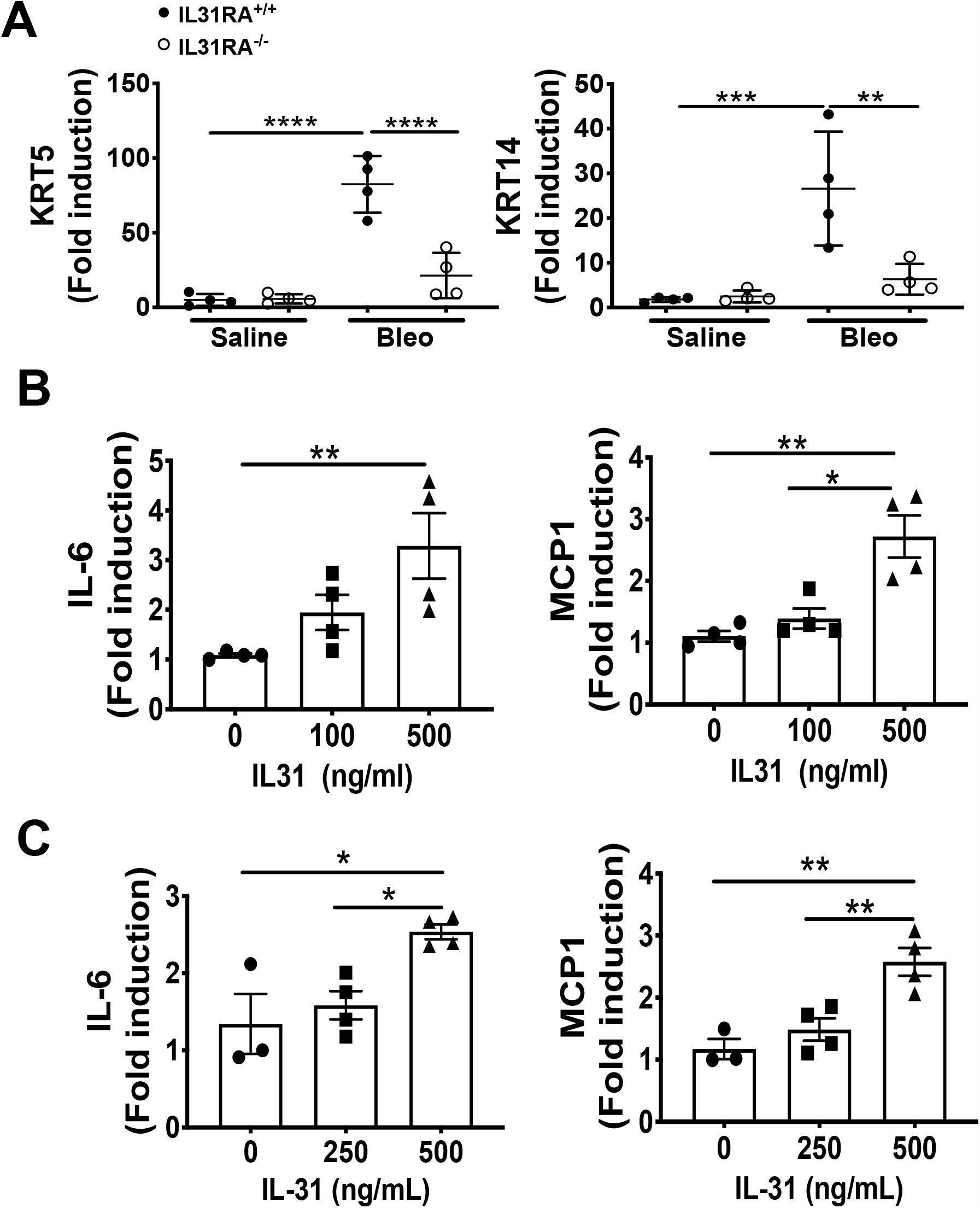
The loss of IL-31RA signaling alters the expression of epithelial cell-specific gene expression during bleomycin-induced pulmonary fibrosis. (A) Reduced expression of epithelial cell-specific genes including KRT5 and KRT14 in the lungs during bleomycin-induced pulmonary fibrosis. (B) Bronchial epithelial cell line BEAS-2B were stimulated with recombinant IL-31 for 24 hours and the transcripts of MCP1 and IL-6 measured using RT-PCR. (C) primary human bronchial epithelial (NHBEC) cells were stimulated with recombinant IL-31 for 24 hours and the transcripts of MCP1 and IL-6 measured using RT-PCR. Data presented as Mean ± SEM (n=4 mice/group). Statistical analysis was performed using one-way ANOVA with Tukey’s multiple comparisons test * p<0.05; ** p<0.01; *** p< 0.005; **** p<0.001.

### Increased frequency of IL-31 producing cells in IPF patients

To determine whether IL-31 is involved in human IPF, IL-31 staining was performed in the lung sections obtained from healthy controls and IPF patients. In contrast to normal lungs, we observed prominent immunostaining for IL-31 in cells populated in the thickened parenchymal areas of IPF lungs (Figure 5A). We quantified the number of IL-31-positive cells and total cells in lung images to assess the accumulation of IL-31-positive cells in IPF lungs compared to healthy lungs. There was a significant increase in the percent of IL-31-positive cells in the lungs from IPF patients compared with healthy controls (Figure 5B). Previous studies have described type 2 CD4 T cells as the main source of IL-31 production in chronic allergic diseases such as atopic dermatitis [11; 38]. To determine the frequency of IL-31 producing CD4 T cells in IPF, PBMCs were stimulated with PMA-ionomycin or medium for 6 h and foxed. Cells were fluorescently stained with IL-31 and IL-4 and analyzed by flow cytometry with a gating strategy as illustrated in Supplementary Figure 2. Notably, the majority of IL-31 producing cells in PBMCs were T cells. CD4 T cells producing IL-31, or IL-4 and IL-31, were significantly increased in PBMCs of IPF patients compared to healthy controls (Figure 5D & 5E).

**Figure 5:**
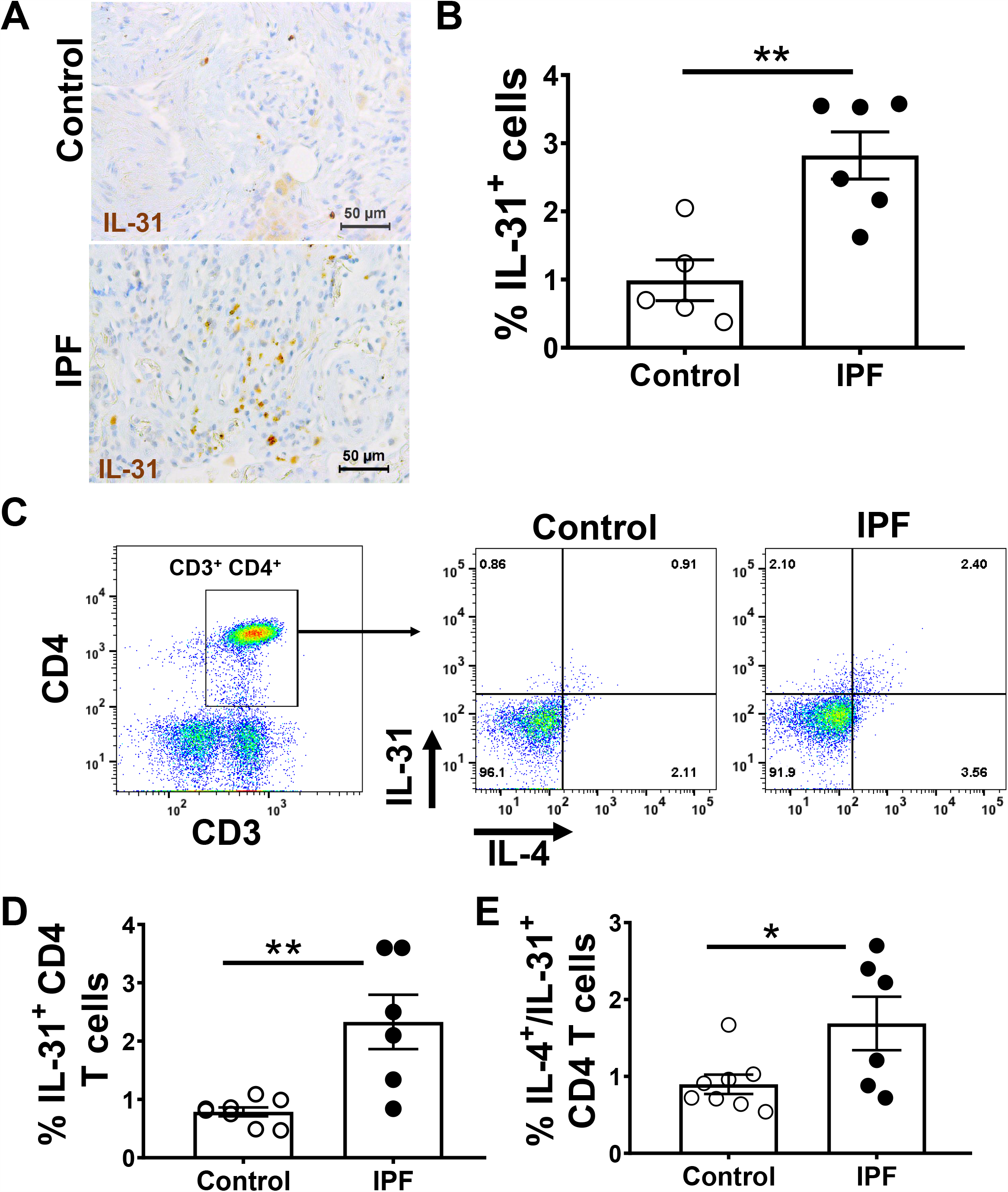
Increased staining and frequency of IL-31-positive CD4 T cells in IPF. Lung sections from IPF and non-IPF control subjects were probed to detect the expression of IL-31 by immunohistochemistry, (A) Representative images of IL-31 expression in non-IPF (top) and IPF (bottom) lung sections; (B) Percent of IL-31^+^ cells quantified in IHC stained lung sections. Data shown as mean ± SEM., ** p<0.01. Peripheral blood mononuclear cells (PBMC) of IPF patients (n=7) and healthy control subjects (n=8) were isolated and stimulated with PMA and ionomycin, intracellular cytokine expression of IL-31 and IL-4 were assessed by flow cytometry. (C) Representative images of CD3 CD4 cells expressing IL-31 and IL-4,unstimulated (top) or stimulated cells with PMA(bottom) from Non-IPF and IPF patients are shown ; (D) Percent of IL-31 and (E) IL4/IL-31-producing CD4 T cells are shown. Data presented as mean ± SEM with n=6-9, with * p<0.05, ** p<0.01. Student *t*-test was used compare the mean between groups.

## Discussion

In this study, we have demonstrated that bleomycin-induced pulmonary fibrosis is dependent on IL-31RA-driven signaling. The loss of IL-31RA signaling was associated with a decrease in hydroxyproline and downregulation of several pro-fibrotic gene transcripts. In addition, our findings demonstrate that the loss of IL-31RA attenuates lung function decline observed during bleomycin induced-pulmonary fibrosis. We also showed that IL-31-positive cells are increased in fibrotic lung lesions of IPF patients and IL-31 is predominantly produced by CD4 T cells. These findings illustrate the potential utility of targeting the IL-31/IL-31RA axis to treat IPF.

The previous studies have identified important roles for Th2 T cell-derived IL-4 and/or IL-13 in the development of pulmonary fibrosis and other fibrotic diseases. However, the role of IL-31/IL-31RA signaling in pulmonary fibrosis has remained unexplored. Our previous study and other research have shown that the administration of exogenous IL-31, or overexpression of IL-31, in transgenic mice induces skin thickening and alters the skin barrier. These findings suggest that IL-31/IL-31RA signaling plays an integral role in inflammation and fibrosis [10; 11]. Previous studies have suggested a potential role for IL-31-driven signaling in the development of fibrosis in various organs including liver [39] and intestine [40]. Shish and colleagues reported increased expression of IL-31RA on intestinal myofibroblasts following treatment with Tl1a, a TNF superfamily 15 protein, and established an association between IL-31RA signaling and colonic fibrosis [40]. Here, we dissected the pro-fibrotic role of the IL-31/IL-31RA axis using a mouse model of bleomycin-induced pulmonary fibrosis. Our studies conducted using IL-31RA knockout mice revealed a significant decrease in hydroxyproline and subsequent improvement in lung function. In addition, whole lung transcriptome analysis revealed a decrease in the expression of ECM-related genes, which supports the role of IL-31/IL-31RA in lung fibrosis. Our findings align with a recent publication demonstrating that in scleroderma IL-31 contributes to skin fibrosis; in this study, a sub-cutaneous mini-pump was implanted in the skin to deliver IL-31 [41]. Together, these studies indicate that IL-31-driven signaling is a key factor driving pulmonary fibrosis. However, additional studies are needed to identify specific mechanisms by which IL-31 contributes to ECM production and decline in lung function during bleomycin-induced pulmonary fibrosis.

Previous studies have identified CD4 helper T cells as a source of IL-31 [11; 12] which is consistent with our findings. We demonstrated a significant increase in IL-31 and IL-4/IL-31-producing CD4^+^ T cells in PBMCs of IPF patients compared with control healthy subjects. The expression of IL-31 was significantly increased in lung tissues of IPF patients compared to control subjects. These findings may indicate that during pulmonary fibrosis development, the TH2 CD4^+^ T cell population contributes to the increased expression of IL-31 in fibrotic lung lesions; within these lung lesions, IL-31 interacts with the unique receptor IL-31RA that is expressed in stromal cells. Collectively, these observations make a compelling case that hematopoietic cell-derived IL-31 might play a role in the pathogenesis of pulmonary fibrosis by activating non-hematopoietic cells such as epithelial cells and mesenchymal cells. Indeed, stimulation of airway epithelial cells with IL-31 induced an increased production of MCP1 and IL-6 cytokines known to play a role in the development of pulmonary fibrosis [42].

We applied the whole lung transcriptome through RNA seq analysis to identify IL-31/IL-31RA -dependent genes and pathways that might participate in IL-31-driven pulmonary fibrosis. Expression of genes associated with the ECM formation was significantly downregulated in IL-31RA^-/-^ mice treated with bleomycin when compared to wildtype mice. Notably, genes associated with epithelial tissue integrity (*Krt5* and *Krt14*), protein Gα signaling, as well as previously reported genes associated with lung function and remodeling (*Arg1, Ccl28* and *Rgs4*) were down regulated in IL-31RA^-/-^ fibrotic mice [43; 44; 45]. Weathington and colleagues have recently reported an association between increased expression of the genes *KCNJ2* and *KRT18* in the setting of asthma, suggesting a strong relationship between lung function and tissue remodeling [46]. In support, genes related to membrane potential regulation, including genes for potassium channels *Kcna2, Kcnh4*, and *Kcnu1*, were upregulated in IL-31RA^-/-^ mice compared to wildtype mice. These findings support a mechanistic pathway that mediates improved lung function in the absence of IL-31RA signaling. Additional investigation is required to explore whether IL-31/IL-31RA signaling contributes to the regulation of potassium channel activity and whether altered keratin gene expression in epithelial cells could mediate collagen deposition and lung function.

In conclusion, our data indicate genetic deletion of IL-31RA significantly attenuates collagen deposition and lung function decline induced during repetitive bleomycin-induced injury and pulmonary fibrosis. In recent years, several cytokines have been implicated in the pathogenesis of pulmonary fibrosis. While IL-31 has been identified in several inflammatory and remodeling diseases, mechanistic studies evaluating IL-31 producing cells in IPF, and the role of IL-31 in pulmonary fibrosis have been missing. Our results provide new evidence that supports the pathogenic role of IL-31RA-driven signaling in bleomycin-induced pulmonary fibrosis. The IL-31-IL31RA axis may serve as a novel therapeutic target for the treatment of pulmonary fibrosis.

## Author Contributions

SKM conceived and designed the research. DJY, and SKM performed the experiments and wrote the manuscript. AGJ performed bioinformatic analysis and edited manuscript. NG provided human IPF samples and edited manuscript.

## Acknowledgments

The authors thank Dr. Brijendra Singh, Dr. Edukulla Ramakrishna, and the veterinary services at Cincinnati Children’s Hospital Medical Center for help in this study.

## Data Availability Statement

Publicly available datasets were analyzed in this study. This data can be found here: https://www.ncbi.nlm.nih.gov/geo/query/acc.cgi?acc=GSE53845

Supplementary Table 1. List of primers used for RT-PCR

Supplementary Table 2: List of IL-31RA-dependent genes

