## Supplemental Data for "The loss of IL-31 signaling attenuates bleomycin-induced pulmonary fibrosis"

***Supplementary Material***

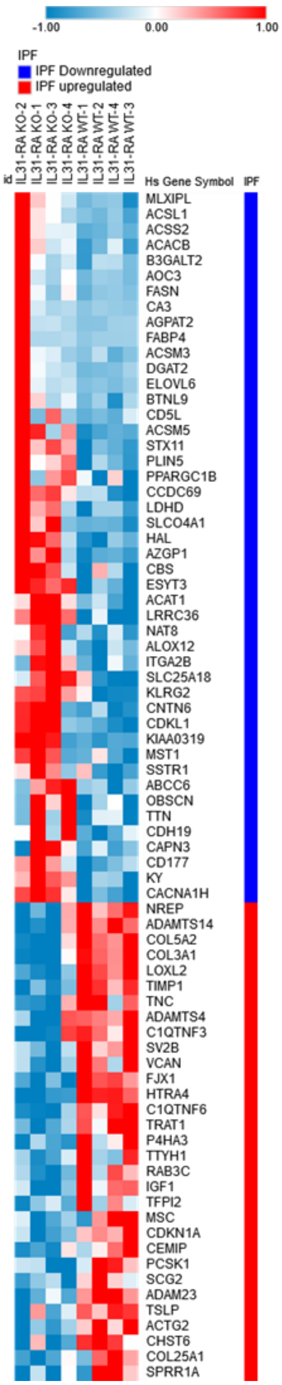


**Supplementary Figure 1.** Heatmap representation of total 77 genes that correlated between IPF and the knockdown of IL-31RA during bleomycin-induced pulmonary fibrosis.

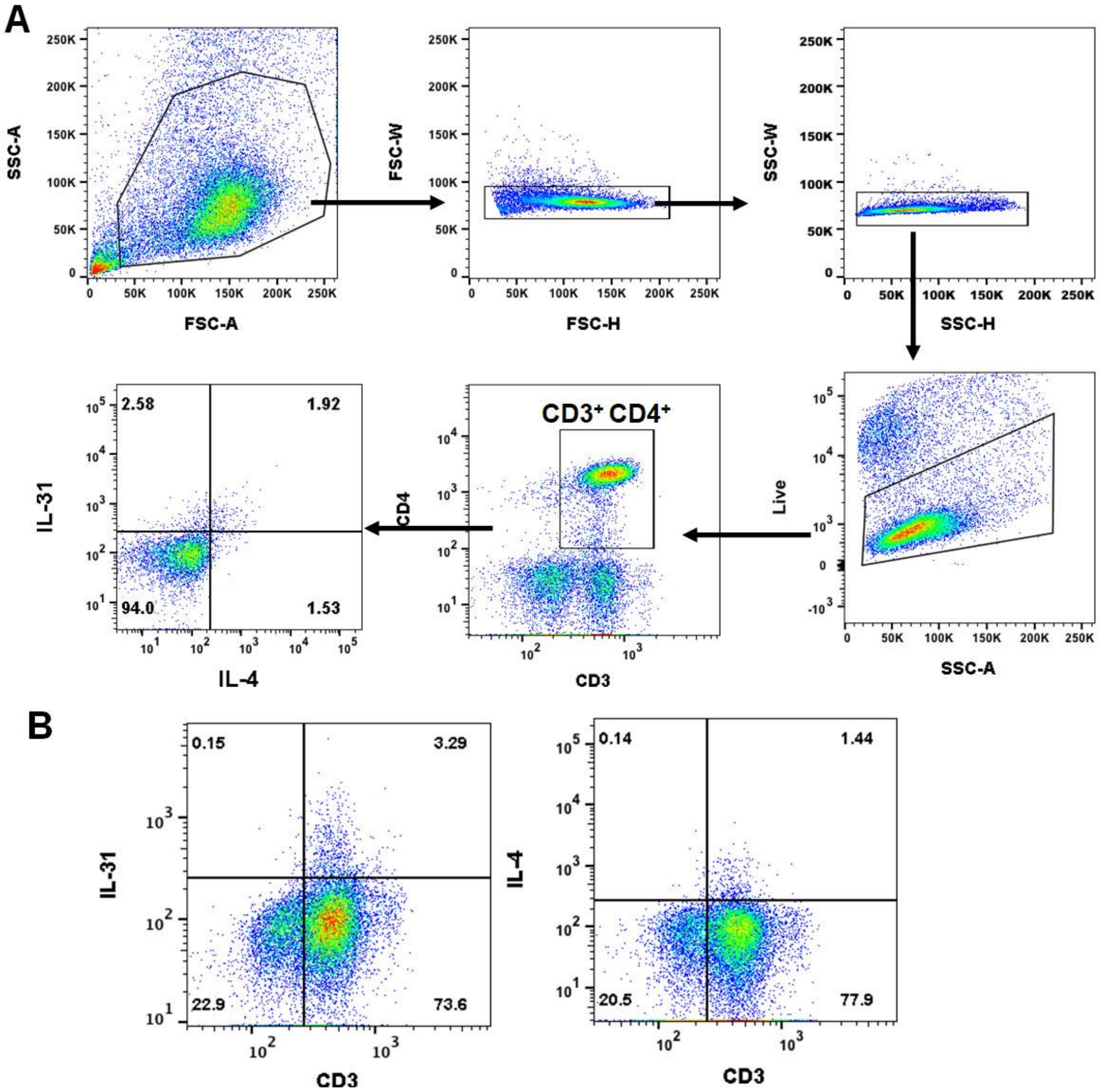


**Supplementary Figure 2. Flow cytometry analysis of IL-31-positive T cells in PBMCs.** (A) Gating strategy used to characterize IL-31-positive cells in PBMC from IPF and healthy subjects. (B) The representative images showing CD3 T cells as a major cell type that produce IL-31 or IL-4 in PBMCs of IPF patients.

**Supplementary Table 1.** List of primers used for RT-PCR.

| Gene ID | Forward primer | Reverse primer |
|---|---|---|
| **mIL6** | TCCAGTTGCCTTCTTGGGAC | GTGTAATTAAGCCTCCGACTTG |
| **HRPT** | GCCCTTGACTATAATGAGTACTTCAGG | TTCAACTTGCGCTCATCTTAGG |
| **mCol1a1** | AGACATGTTCAGCTTTGTGGAC | GCAGCTGACTTCAGGGATG |
| **mCol3a1** | TCCCCTGGAATCTGTGAATC | TGAGTCGAATTGGGGAGAAT |
| **mFn1** | CGGAGAGAGTGCCCCTACTA | CGATATTGGTGAATCGCAGA |
| **mMMP-13** | CTGGCACACGCTTTTCCTCCTG | GGGCTGGGTCACACTTCTCTGGT |
| **mTimp1** | GCAAAGAGCTTTCTCAAAGACC | AGGGATAGATAAACAGGGAAACACT |
| **mKrt5** | CAGAGCTGAGGAACATGCAG | CATTCTCAGCCGTGGTACG |
| **mKrt14** | ATCGAGGACCTGAAGAGCAA | TCGATCTGCAGGAGGACATT |
| **hMCP1** | GCTCATAGCAGCCACCTTCA | ACAATGGTCTTGAAGATCACAGC |
| **hIL-6** | CAGGAGCCCAGCTATGAACT | GAAGGCAGCAGGCAACAC |

**Supplementary Table 2.** The list of genes differentially expressed in IL-31RA knockout mice during bleomycin-induced pulmonary fibrosis.

| Genes Upregulated in IL-31RA KO Mice | | | Genes Downregulated in IL-31RA KO Mice | | |
|---|---|---|---|---|---|
| **Gene Symbol** | **Log2 Fold** | **p-Val** | | **Log2 Fold** | **p-Val** |
| Chil4 | 2.673613 | 0.000142 | Sox18 | -0.58571 | 6.6E-08 |
| 4933400F21Rik | 2.449708 | 0.002679 | Psapl1 | -0.59137 | 0.013305 |
| A330074K22Rik | 2.196978 | 0.000553 | Gjb5 | -0.59174 | 0.005489 |
| AI593442 | 1.968621 | 0.000628 | Col4a2 | -0.59288 | 8.66E-10 |
| Clca1 | 1.909314 | 4.68E-10 | Carmn | -0.59383 | 0.023858 |
| 5830411N06Rik | 1.822686 | 0.00576 | Nrcam | -0.59543 | 3.96E-06 |
| Krt79 | 1.752798 | 1.81E-06 | Tmem45a | -0.59679 | 6.92E-08 |
| Fabp12 | 1.731421 | 0.001595 | Adamts2 | -0.59682 | 1.01E-07 |
| Olfr750 | 1.717942 | 0.026154 | Ccl2 | -0.59743 | 0.000452 |
| Adgrf3 | 1.591881 | 0.010797 | Id3 | -0.59908 | 3.42E-12 |
| Leap2 | 1.569674 | 0.027845 | Pcdhb19 | -0.59999 | 0.016732 |
| Sult2a1 | 1.553427 | 0.049406 | Fhl2 | -0.60137 | 0.000547 |
| Tmem178b | 1.529699 | 0.004423 | Csmd1 | -0.60385 | 0.017902 |
| Hsd17b13 | 1.525838 | 0.029312 | Klhdc8a | -0.60541 | 4.06E-06 |
| Ngp | 1.522654 | 0.02251 | Chodl | -0.60661 | 0.004555 |
| Lctl | 1.519688 | 0.036391 | Lhx6 | -0.60747 | 2.73E-05 |
| Mir7013 | 1.517201 | 0.011605 | Bmper | -0.6082 | 3.86E-05 |

| | | | | | |
|---|---|---|---|---|---|
| Fcgbp | 1.509645 | 1.37E-08 | Prss35 | -0.61025 | 0.004394 |
| Cd300e | 1.506774 | 9.07E-08 | Alox15 | -0.6106 | 0.035185 |
| Cyp1a1 | 1.477192 | 5.82E-05 | Apcdd1 | -0.61292 | 2.27E-05 |
| Pkdrej | 1.472519 | 0.00013 | Galnt16 | -0.61366 | 0.000523 |
| Slc6a20a | 1.471947 | 1.48E-11 | Ccdc80 | -0.6146 | 1.98E-08 |
| Ceacam10 | 1.455468 | 0.015381 | Nrg1 | -0.61622 | 1.43E-05 |
| Ces2c | 1.437188 | 0.049946 | Fstl1 | -0.61667 | 3.06E-10 |
| Hist1h4h | 1.409537 | 1.33E-07 | Tnfrsf12a | -0.6173 | 5.58E-07 |
| 4930433N12Rik | 1.394948 | 0.006936 | P2ry14 | -0.61762 | 2.28E-08 |
| Colq | 1.390389 | 4.14E-05 | Wscd2 | -0.61825 | 0.004752 |
| C730036E19Rik | 1.37155 | 0.001619 | Disp2 | -0.61923 | 4.36E-09 |
| Entpd8 | 1.288417 | 0.00099 | Baalc | -0.61984 | 0.027398 |
| Ccl20 | 1.28541 | 0.010986 | Adgrb2 | -0.62059 | 0.020691 |
| Rpph1 | 1.284619 | 0.016616 | Alpl | -0.62141 | 0.002383 |
| Art2b | 1.265003 | 0.022463 | Col5a2 | -0.62162 | 3.05E-08 |
| Gm15713 | 1.25535 | 0.034382 | Nxpe5 | -0.62429 | 0.003603 |
| Gm16863 | 1.254123 | 0.041035 | Junb | -0.6263 | 1.63E-06 |
| Tbata | 1.239877 | 0.000767 | Tnc | -0.62762 | 1.92E-05 |
| Cst8 | 1.225231 | 0.023483 | Sfrp1 | -0.6279 | 8.35E-09 |
| Muc5ac | 1.215486 | 7.82E-11 | Aplnr | -0.62936 | 0.000942 |
| 9230117E06Rik | 1.212656 | 0.029542 | Adamts4 | -0.63026 | 7.53E-09 |

| | | | | | |
|---|---|---|---|---|---|
| Lrat | 1.212301 | 2.85E-09 | Mt1 | -0.63152 | 0.00417 |
| Gria1 | 1.209916 | 1.2E-05 | Rgs4 | -0.6325 | 2.11E-06 |
| Slc4a1 | 1.20287 | 0.004447 | Acta2 | -0.63381 | 6.64E-05 |
| Tecrl | 1.201598 | 1.15E-06 | Serpinb9b | -0.63412 | 3.59E-05 |
| Gypa | 1.192801 | 0.010383 | Sema7a | -0.63563 | 5.2E-07 |
| Slc7a10 | 1.186105 | 1.28E-08 | Ramp3 | -0.63749 | 0.00338 |
| Stac | 1.18021 | 0.046521 | Gm13889 | -0.6394 | 3.26E-07 |
| Gbp10 | 1.179485 | 0.004405 | S100a14 | -0.6394 | 3.29E-09 |
| Gpr33 | 1.164263 | 0.007738 | Fkbp5 | -0.6421 | 0.000412 |
| Btnl9 | 1.149575 | 0.012233 | Col1a1 | -0.64445 | 6.06E-06 |
| Kcnu1 | 1.144672 | 0.001337 | Aspn | -0.64457 | 7.53E-10 |
| Grip2 | 1.142652 | 0.006848 | Mark1 | -0.64586 | 7.1E-08 |
| Gm29508 | 1.132961 | 0.027103 | Shf | -0.64659 | 0.009229 |
| Acoxl | 1.125048 | 2.95E-13 | Rtn4rl2 | -0.64811 | 0.028092 |
| Rmrp | 1.120263 | 0.023983 | Fam181b | -0.64979 | 0.005833 |
| Gjb1 | 1.0942 | 0.006782 | Cers3 | -0.6533 | 0.000359 |
| Lipf | 1.092653 | 1.41E-05 | Il20rb | -0.65552 | 0.000226 |
| Cdh16 | 1.09093 | 0.006542 | 2200002D01Rik | -0.6596 | 9.24E-05 |
| Ifi214 | 1.089664 | 0.003476 | Actg2 | -0.65998 | 0.00106 |
| Adgre4 | 1.087422 | 3.41E-05 | Bhlhe22 | -0.66353 | 0.003501 |
| Gm15441 | 1.085195 | 0.041732 | Aldh1a3 | -0.66583 | 0.003011 |

| | | | | | |
|---|---|---|---|---|---|
| Bex2 | 1.081454 | 0.003401 | Mboat2 | -0.66797 | 5.34E-07 |
| Cecr6 | 1.078146 | 0.024324 | Serpina3n | -0.66852 | 2.33E-06 |
| Gc | 1.070781 | 0.029874 | Fgf2 | -0.67303 | 0.000454 |
| Aass | 1.070057 | 1.53E-08 | Bean1 | -0.67317 | 0.005458 |
| Nrn1 | 1.061699 | 2.14E-07 | Ptx3 | -0.67402 | 0.01925 |
| Slc1a2 | 1.047459 | 9.76E-05 | 1500015O10Rik | -0.67547 | 0.022997 |
| Apol11b | 1.041322 | 0.008488 | Clec10a | -0.68523 | 7.38E-05 |
| A930006K02Rik | 1.039347 | 0.000676 | Vwa1 | -0.68636 | 1.89E-06 |
| Ntn5 | 1.037847 | 0.01952 | P4ha3 | -0.68752 | 0.00017 |
| Sbk2 | 1.030719 | 0.018273 | Tnfrsf11b | -0.6895 | 2.14E-05 |
| Slfn5os | 1.029082 | 0.030258 | Wisp1 | -0.69371 | 6.3E-06 |
| 6330407A03Rik | 1.028617 | 0.037364 | Arg1 | -0.6942 | 0.008545 |
| Cd177 | 1.015409 | 0.020052 | Gper1 | -0.69561 | 0.003943 |
| S1pr5 | 1.009543 | 1.04E-05 | Cnn1 | -0.69567 | 0.003332 |
| Hc | 1.006359 | 1.3E-21 | Arc | -0.69687 | 0.0076 |
| Gjb6 | 1.004205 | 2.01E-06 | Ms4a4a | -0.7054 | 6.87E-05 |
| 1700128F08Rik | 1.00216 | 0.00939 | Rtn4r | -0.7085 | 0.046516 |
| Snca | 0.994158 | 2.63E-05 | Ceacam16 | -0.70922 | 0.027628 |
| Ubd | 0.983973 | 0.042134 | Tubb3 | -0.71689 | 0.003071 |
| Gm14085 | 0.981087 | 0.01794 | Mgp | -0.71866 | 4.23E-13 |
| Tat | 0.966249 | 0.036146 | Ptgdr2 | -0.72169 | 0.038499 |

| | | | | | |
|---|---|---|---|---|---|
| Hmgcs2 | 0.965869 | 2.64E-08 | Tagln | -0.72528 | 3.43E-08 |
| Cxcr1 | 0.952686 | 0.000319 | Stc2 | -0.72709 | 0.001155 |
| Tex11 | 0.952675 | 2.06E-08 | Dbn1 | -0.73075 | 8.1E-09 |
| Heatr9 | 0.952508 | 0.026835 | Saa3 | -0.73137 | 0.019986 |
| Capn9 | 0.951696 | 0.040205 | Tubb2b | -0.73241 | 0.00177 |
| Avpr1a | 0.942627 | 0.009621 | Mycn | -0.73252 | 0.020154 |
| Cntn4 | 0.9392 | 0.003044 | Wnk3 | -0.7364 | 0.031459 |
| Pigr | 0.938587 | 1.45E-16 | Flnc | -0.74101 | 7.69E-05 |
| Itih1 | 0.925434 | 0.047826 | Clrn1 | -0.74147 | 0.041874 |
| Ppbp | 0.920734 | 3.31E-06 | Syt13 | -0.74749 | 0.024204 |
| Ccr3 | 0.920423 | 0.03171 | Rnf152 | -0.74784 | 2.03E-05 |
| Treml4 | 0.920215 | 1.7E-05 | Cldn4 | -0.7508 | 1.37E-08 |
| Cyp26b1 | 0.918901 | 0.047801 | Fmod | -0.75119 | 3E-08 |
| Itih4 | 0.915934 | 8.47E-18 | Ltbp2 | -0.75761 | 2.11E-09 |
| 4930512B01Rik | 0.909073 | 0.030949 | Mnd1 | -0.75825 | 0.001818 |
| Vnn3 | 0.90534 | 0.000768 | Hoxd8 | -0.76167 | 0.01932 |
| 4931406H21Rik | 0.902503 | 0.006289 | Gal3st2 | -0.76357 | 0.015301 |
| Lrrc17 | 0.90239 | 0.000154 | Igsf1 | -0.7699 | 0.011721 |
| Cd8a | 0.901348 | 0.00126 | Inhba | -0.77317 | 2.03E-11 |
| Ctcfl | 0.899915 | 0.005459 | Gdf6 | -0.77325 | 0.000578 |
| Klre1 | 0.895238 | 0.000822 | Shc4 | -0.77344 | 0.000206 |

| | | | | | |
|---|---|---|---|---|---|
| Awat2 | 0.885709 | 0.025555 | Cd248 | -0.77527 | 6.91E-13 |
| Proz | 0.867426 | 0.001534 | Rasl10b | -0.78815 | 0.00022 |
| Bpifa2 | 0.865212 | 0.003172 | Fn1 | -0.78956 | 1.17E-11 |
| Chil1 | 0.863091 | 2.97E-25 | Xirp2 | -0.78966 | 0.005784 |
| Acod1 | 0.863042 | 0.010323 | Pcdhgb8 | -0.79073 | 0.013895 |
| Ros1 | 0.86182 | 0.000668 | Scn8a | -0.79558 | 0.015527 |
| Gm2115 | 0.860746 | 0.003023 | Snord47 | -0.80094 | 0.010658 |
| Cd163l1 | 0.85727 | 0.004771 | Dkk2 | -0.8121 | 0.000905 |
| Hpcal4 | 0.854495 | 0.000294 | Tram1l1 | -0.81561 | 0.005958 |
| Fxyd7 | 0.853736 | 0.029262 | Mmp13 | -0.82059 | 7.79E-06 |
| Cth | 0.851512 | 2.17E-05 | Tmem252 | -0.82195 | 0.001738 |
| Gm6654 | 0.848027 | 0.014921 | Dio2 | -0.82808 | 0.000365 |
| Mgam | 0.845967 | 0.026191 | Smpd3 | -0.83053 | 1.48E-06 |
| Slc13a4 | 0.830348 | 0.00078 | C130080G10Rik | -0.83197 | 0.012641 |
| Slfn14 | 0.822254 | 0.04703 | Col28a1 | -0.83301 | 5.96E-10 |
| Hkdc1 | 0.820164 | 0.00248 | Lox | -0.83378 | 3.78E-14 |
| Kcnh4 | 0.818558 | 0.032971 | Prg4 | -0.84001 | 6.89E-05 |
| Phf24 | 0.818176 | 1.96E-05 | Aicda | -0.84513 | 0.031228 |
| Galnt13 | 0.817111 | 0.000362 | Ackr1 | -0.84516 | 0.001684 |
| A930001A20Rik | 0.815555 | 0.013087 | Iglon5 | -0.84796 | 0.009439 |
| Mbl1 | 0.80788 | 0.021332 | Syt8 | -0.85556 | 0.022212 |

| | | | | | |
|---|---|---|---|---|---|
| Tff2 | 0.805843 | 2.28E-05 | Il22ra2 | -0.8665 | 0.001877 |
| Nalcn | 0.802981 | 0.010329 | Kcnh6 | -0.86707 | 0.040281 |
| Mme | 0.802494 | 4.09E-14 | Kcnj10 | -0.87587 | 0.003001 |
| Dio1 | 0.800167 | 0.009147 | Fcna | -0.87819 | 0.000291 |
| Pbld2 | 0.798502 | 4.33E-05 | Prss22 | -0.8832 | 0.007636 |
| Cyp2ab1 | 0.797159 | 0.006177 | Gpr176 | -0.89271 | 2.73E-06 |
| Bcan | 0.796814 | 0.010578 | Pappa | -0.8938 | 1.47E-07 |
| Sh2d1a | 0.791745 | 0.012118 | Ccl12 | -0.89815 | 2.82E-05 |
| Dnah7c | 0.790012 | 0.009572 | Cd5l | -0.89933 | 0.033353 |
| Grik4 | 0.788655 | 0.031628 | Htra4 | -0.90532 | 4.16E-07 |
| Vipr2 | 0.786055 | 1.35E-07 | Padi1 | -0.90562 | 0.037523 |
| Lmntd1 | 0.784217 | 0.014322 | Snai1 | -0.90715 | 5.46E-13 |
| Gm20257 | 0.7818 | 0.001175 | Prrx2 | -0.90869 | 0.032842 |
| D030025P21Rik | 0.772405 | 0.029944 | Calcr | -0.90955 | 0.036381 |
| Pisd-ps1 | 0.771626 | 2.15E-15 | Eln | -0.91772 | 3.64E-13 |
| Ptgs2os2 | 0.771197 | 0.000237 | Slc17a2 | -0.92535 | 0.039815 |
| Bend6 | 0.767975 | 0.002298 | Xcl1 | -0.92751 | 0.020171 |
| Ppp2r2c | 0.762647 | 0.002091 | Wfdc12 | -0.96427 | 0.000174 |
| Arhgef38 | 0.758907 | 0.005284 | Timp1 | -0.96694 | 3.14E-11 |
| Tubb1 | 0.751273 | 0.000656 | Spock3 | -0.97767 | 0.03904 |
| Rgs7 | 0.750917 | 0.046371 | Zbtb16 | -0.98012 | 4.34E-08 |

| | | | | | |
|---|---|---|---|---|---|
| Srpk3 | 0.750212 | 0.00254 | Ptk6 | -0.98222 | 0.000669 |
| 1810044D09Rik | 0.747929 | 0.027686 | Mapk4 | -0.99586 | 0.00122 |
| Cd8b1 | 0.745079 | 0.020092 | Fbln2 | -0.99785 | 2.95E-23 |
| Gm32014 | 0.743705 | 0.028476 | Il6 | -0.99836 | 0.036906 |
| Plppr4 | 0.74287 | 0.026068 | Mest | -1.0017 | 1.93E-11 |
| Agtr1b | 0.742029 | 0.033589 | Olfr558 | -1.00651 | 0.025765 |
| 2310040G24Rik | 0.734871 | 0.008535 | Tceal3 | -1.00911 | 0.045607 |
| 1110020A21Rik | 0.733519 | 0.016566 | Mmp10 | -1.01046 | 0.009625 |
| Samd15 | 0.733279 | 0.024111 | Nts | -1.02542 | 0.003267 |
| Hepacam | 0.730381 | 0.022396 | Nrxn1 | -1.03011 | 1.44E-11 |
| Krt23 | 0.729252 | 1.1E-06 | Serpina3m | -1.03079 | 0.002163 |
| Snora81 | 0.728818 | 0.033893 | Pla1a | -1.03092 | 3.5E-14 |
| Marco | 0.726037 | 2.01E-07 | Rnf165 | -1.03457 | 0.006203 |
| Crispld1 | 0.723924 | 0.003837 | Cd300c | -1.03803 | 0.023686 |
| Pisd-ps2 | 0.711691 | 1.9E-10 | Nr1h4 | -1.03811 | 0.028219 |
| Gpr18 | 0.711572 | 0.003812 | Resp18 | -1.03899 | 0.003886 |
| Acot1 | 0.706935 | 7.47E-06 | Rgs16 | -1.03921 | 2.41E-09 |
| Cntfr | 0.704008 | 0.0114 | Hspb7 | -1.06488 | 6.62E-06 |
| S100g | 0.703372 | 1.07E-09 | Rgs5 | -1.06665 | 8.08E-28 |
| Nrg2 | 0.697571 | 0.004323 | Synpo2l | -1.07442 | 0.001955 |
| 1810041L15Rik | 0.695114 | 0.007446 | Gjb4 | -1.0857 | 0.010558 |

| | | | | | |
|---|---|---|---|---|---|
| Pcdhga4 | 0.695099 | 0.00405 | Olfr78 | -1.09502 | 0.00636 |
| Sgpp2 | 0.694221 | 2.4E-15 | Tarm1 | -1.10022 | 0.021312 |
| Tlr11 | 0.692812 | 0.019532 | Ereg | -1.10422 | 0.000217 |
| Gm20597 | 0.692353 | 0.045531 | A2m | -1.10641 | 0.016 |
| Myo15 | 0.69213 | 0.009936 | Egfem1 | -1.11408 | 3.23E-07 |
| Ar | 0.691972 | 4.18E-05 | Prtn3 | -1.1192 | 0.026912 |
| Kcna2 | 0.68989 | 0.000989 | Tfap2a | -1.12856 | 4E-05 |
| Cttnbp2 | 0.689289 | 0.00124 | 3300005D01Rik | -1.13515 | 0.007248 |
| Idi1 | 0.688066 | 3.36E-11 | Mchr1 | -1.13824 | 0.006389 |
| Spta1 | 0.685678 | 0.022097 | Slc24a1 | -1.14122 | 0.026726 |
| B230217C12Rik | 0.68486 | 0.011896 | Tpsb2 | -1.14391 | 0.04054 |
| Esrrg | 0.683934 | 0.000145 | 2210408I21Rik | -1.14567 | 3.64E-05 |
| Lamp3 | 0.681293 | 5.48E-18 | Fabp7 | -1.15544 | 0.000257 |
| Slc22a3 | 0.67869 | 0.03921 | Dspp | -1.15652 | 0.001355 |
| Aadac | 0.675362 | 0.011606 | Mt2 | -1.1627 | 1.64E-06 |
| Ces1f | 0.674206 | 0.002186 | Peg3 | -1.17064 | 1.59E-16 |
| 9330159F19Rik | 0.673698 | 0.001874 | Dio3os | -1.18736 | 0.013563 |
| Dpp6 | 0.670869 | 0.002994 | Serpinb2 | -1.19177 | 0.000283 |
| Edn3 | 0.667307 | 8.1E-05 | Krt5 | -1.1923 | 0.002317 |
| Cpm | 0.666875 | 3.47E-14 | Atp13a5 | -1.19331 | 0.007924 |
| Nr3c2 | 0.665198 | 2.29E-08 | Xirp1 | -1.20639 | 0.037702 |

| | | | | | |
|---|---|---|---|---|---|
| Gm11744 | 0.662299 | 0.026446 | Ankrd34a | -1.20732 | 0.037274 |
| Tcf23 | 0.658968 | 0.010798 | Dok5 | -1.21049 | 0.033988 |
| Esr2 | 0.655701 | 0.003412 | Ankrd34b | -1.21207 | 0.000979 |
| Itk | 0.653062 | 0.001474 | Grem1 | -1.21501 | 0.000758 |
| 2610307P16Rik | 0.652735 | 0.015737 | Cdh3 | -1.23744 | 2.08E-07 |
| Inmt | 0.6498 | 1.21E-06 | Nog | -1.25027 | 2.96E-05 |
| Alas2 | 0.649058 | 0.001107 | Wnt10a | -1.25514 | 7.52E-05 |
| Slc38a5 | 0.648871 | 0.00025 | Angptl7 | -1.26904 | 3.13E-06 |
| Nkg7 | 0.647346 | 0.002451 | Moxd1 | -1.28491 | 1.19E-06 |
| Mfsd2a | 0.645631 | 4.61E-07 | Fst | -1.306 | 2.89E-12 |
| Slco4c1 | 0.64364 | 2.3E-07 | Calcb | -1.3085 | 2.1E-05 |
| Adra2a | 0.643116 | 0.010946 | Chl1 | -1.31269 | 1.83E-15 |
| Lrp2 | 0.637266 | 5.62E-11 | Frzb | -1.31456 | 2.76E-06 |
| 5330413P13Rik | 0.63491 | 0.007508 | 4930546K05Rik | -1.32176 | 0.021605 |
| Snhg11 | 0.633916 | 8.75E-09 | Zfp985 | -1.33229 | 0.000194 |
| Zpbp | 0.630773 | 0.036032 | Rspo4 | -1.36 | 0.036528 |
| Gm12250 | 0.630132 | 1.2E-05 | Pgbd5 | -1.36586 | 0.000363 |
| Gm5084 | 0.630091 | 0.029315 | Krt4 | -1.37121 | 0.012891 |
| Dcdc2a | 0.629278 | 0.000477 | Serpinb5 | -1.3773 | 0.041347 |
| Gpt | 0.62757 | 5.27E-07 | Dio3 | -1.38034 | 1.48E-07 |
| 2010005H15Rik | 0.622056 | 0.039898 | Gcnt4 | -1.40666 | 0.000159 |

| | | | | | |
|---|---|---|---|---|---|
| Colgalt2 | 0.621817 | 0.000869 | Bpifa1 | -1.40944 | 7.56E-12 |
| Sfxn4 | 0.620845 | 0.002212 | Kcnt1 | -1.42272 | 0.011289 |
| Ptpn20 | 0.619916 | 0.006704 | Kng2 | -1.42341 | 1.13E-07 |
| Alox12 | 0.617404 | 0.001851 | Fgf23 | -1.42466 | 0.018705 |
| Wdr95 | 0.617079 | 0.004089 | Kcng2 | -1.46575 | 0.024785 |
| Plin1 | 0.617052 | 0.010172 | Pnma2 | -1.4763 | 0.001024 |
| Egfl6 | 0.61703 | 3.24E-12 | Krt20 | -1.52728 | 0.03796 |
| 2810459M11Rik | 0.614825 | 0.034506 | Gm12709 | -1.55574 | 0.029477 |
| Nipal1 | 0.612598 | 0.00139 | Krt14 | -1.57194 | 0.000799 |
| Gzmb | 0.611966 | 0.03493 | Esm1 | -1.60428 | 4.26E-14 |
| Cd19 | 0.610868 | 0.006056 | Ccl28 | -1.61613 | 9.19E-06 |
| Adra1a | 0.609117 | 0.001359 | Npas3 | -1.61739 | 0.004178 |
| H2-K2 | 0.607889 | 0.007169 | Krt17 | -1.65181 | 0.000267 |
| Pla2g4f | 0.604119 | 0.00071 | C430002N11Rik | -1.66975 | 0.002388 |
| Sftpa1 | 0.602931 | 7.46E-13 | Tmprss11g | -1.6972 | 7.12E-05 |
| Akap14 | 0.594143 | 0.003335 | Tnfsf18 | -1.74337 | 3.9E-05 |
| Faim2 | 0.593782 | 0.041433 | Cwh43 | -1.77127 | 3.31E-05 |
| Plekhd1os | 0.593636 | 0.047726 | Nppa | -1.79786 | 0.004488 |
| Ric3 | 0.592527 | 0.002016 | Calcoco2 | -1.85255 | 0.006209 |
| Cntn1 | 0.589763 | 0.049646 | Cemip | -1.89049 | 1.7E-13 |
| Dhtkd1 | 0.588409 | 0.017916 | Krt6a | -1.94824 | 0.013342 |

| | | | | | |
|---|---|---|---|---|---|
| C530008M17Rik | 0.586904 | 0.012049 | Pkp1 | -2.02817 | 9.69E-06 |
| | | | Tmem267 | -2.78047 | 1.03E-48 |
| | | | Erdr1 | -4.59138 | 6.42E-20 |
